# Mechanical Loading Recovers Bone but not Muscle Lost During Unloading

**DOI:** 10.1101/2020.09.01.275990

**Authors:** Andrew R. Krause, Toni A. Speacht, Jennifer L. Steiner, Charles H. Lang, Henry J. Donahue

**Affiliations:** Department of Orthopaedics, Penn State College of Medicine, Hershey, PA 17033-2391 USA; Department of Nutrition, Food and Exercise Science, Florida State University, Tallahassee, FL 32306 USA; Department of Cellular and Molecular Physiology, Penn State College of Medicine, Hershey, PA 17033-2391 USA; Bone Engineering, Science and Technology (BEST) Lab, Department of Biomedical Engineering, Virginia Commonwealth University College of Engineering, Richmond, VA 23284-3067 USA

**Keywords:** hind limb suspension, re-loading, bone loss, muscle loss

## Abstract

Space travel and prolonged bed rest are examples of mechanical unloading that induce significant muscle and bone loss. To explore interactions between skeletal bone and muscle during reloading, we hypothesized that acute external mechanical loading of bone in combination with re-ambulation facilitates proportional recovery of bone and muscle lost during hind limb suspension (HLS) unloading. Adult male C57Bl/6J mice were assigned to a HLS or time-matched ground control (GC) group. After 2-weeks of HLS, separate groups of mice were studied at day 14 (no re-ambulation), day 28 (14 days re-ambulation) and day 56 (42 days re-ambulation); throughout the re-ambulation period, one limb received mechanical loading and the contralateral limb served as an internal control. HLS induced loss of trabecular bone volume (BV/TV; -51%±2%) and muscle weight (*-*15%±2%) compared to GC at day 14. At day 28, the left tibia (re-ambulation only) of HLS mice had recovered 20% of BV/TV lost during HLS, while the right tibia (re-ambulation and acute external mechanical loading) recovered to GC values of BV/TV (∼100% recovery). At day 56, the right tibia continued to recover bone for some outcomes (trabecular BV/TV, trabecular thickness), while the left limb did not. Cortical bone displayed a delayed response to HLS, with a 10% greater decrease in BV/TV at day 28 compared to day 14. In contrast to bone, acute external mechanical loading during the re-ambulation period did not significantly increase muscle mass or protein synthesis in the gastrocnemius, compared to re-ambulation alone.

## Introduction

Space travel, particularly missions to Mars, will expose astronauts to extended periods of mechanical unloading that produce significant bone and muscle loss [1-5]. While unloading causes a concomitant atrophy of bone and muscle, of equal clinical importance is the subsequent reloading period. Decreased structure and function of the musculoskeletal system associated with osteopenia and sarcopenia following unloading increases risk of injury during re-ambulation [6-9]. Therefore, there is a need to study the reloading period and better understand the bone-muscle relationship during recovery from unloading, and potentially identify new interventions to facilitate recovery of the musculoskeletal system. The reloading period is a relatively under-investigated area in the context of musculoskeletal unloading. Therefore, in this study we evaluated the catabolic effects associated with hind limb suspension (HLS) unloading, followed by re-ambulation in the presence or absence of acute external mechanical loading.

Emerging evidence suggests that changes in bone structure and function, as might occur during unloading or loading, may affect muscle and vice versa [10-13]. For instance, in vitro exposure of osteocytic cells to shear stress, as would occur from mechanical loading, results in release of PGE2 and Wnt3a both of which support myogenesis [14]. These data suggest that loading bone in vivo has the potential to affect myogenesis in vivo. Other bone secreted factors, such as osteocalcin and RANKL, can also impact muscle metabolism [15-17]. Additionally, myokines such as myostatin, secreted by muscle can negatively impact bone remodeling [18, 19]. However, myoblast conditioned media has been shown to be bone anabolic both in vitro and in vivo [20] suggesting the relationship between bone and muscle, especially in the context of loading and unloading, is not well understood. Therefore, we examined the hypothesis that acute external mechanical loading facilitates the proportional recovery of bone and muscle loss resulting from unloading.

## Results

### Trabecular Outcomes in Ground Control Mice

Compared to baseline scans performed on day 0, several trabecular endpoints were decreased in the tibia of GC mice **(Fig 1, white bars)**. Trabecular bone volume fraction (BV/TV) and trabecular thickness decreased whereas trabecular separation increased at day 14, compared to baseline scans (**Fig 1, white bars**); these values did not change further on day 28 or 56, compared to day 14. Additionally, trabecular number, connectivity density and trabecular bone mineral density all decreased at day 14 compared to baseline scans (**Fig 2, white bars**); the catabolic change from baseline for trabecular number and connectivity density were greater at day 56 compared to day 14 (**Fig 2, white bars**).

**Figure 1:**
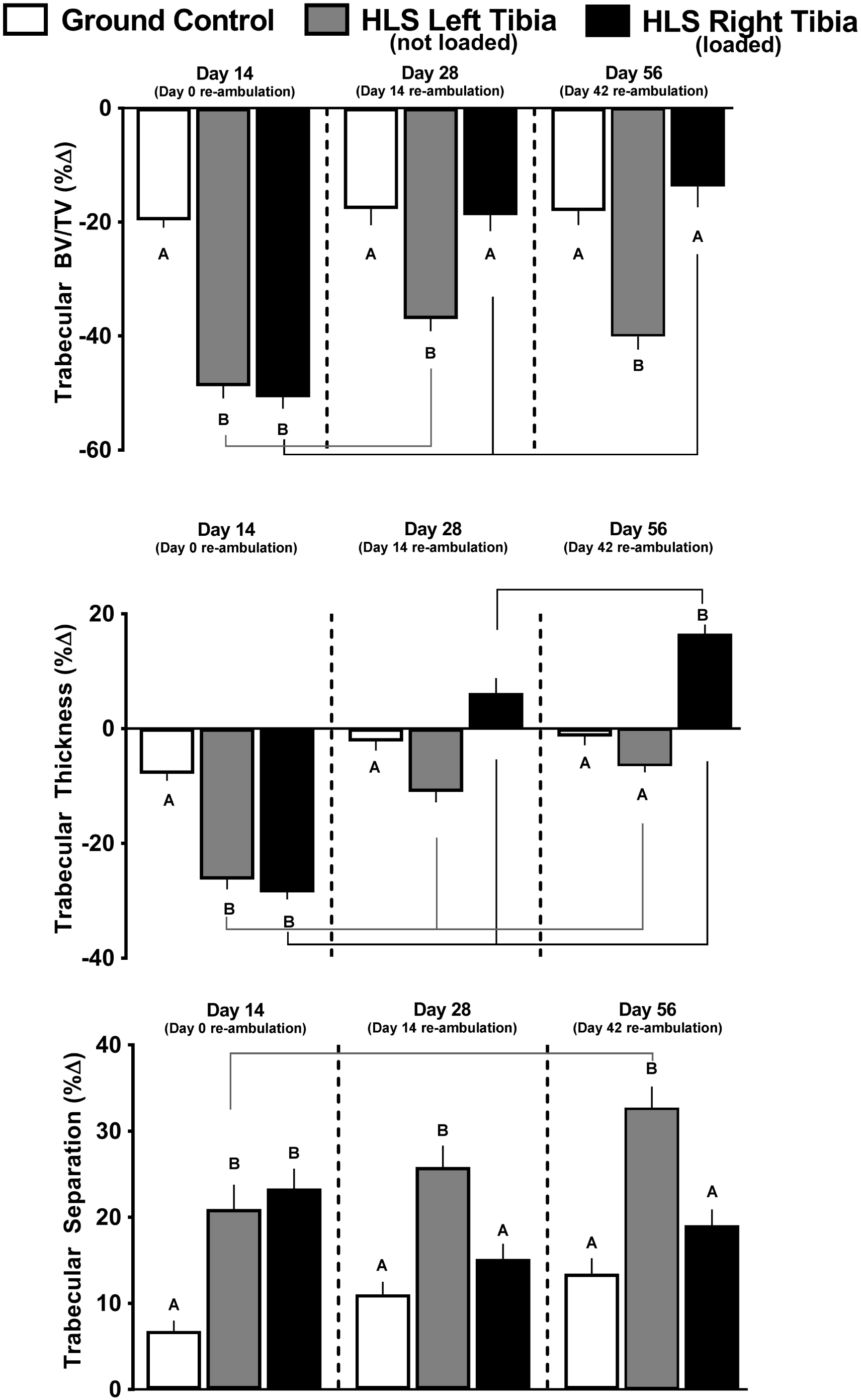
Effect of HLS, re-ambulation only and re-ambulation and mechanical loading on trabecular micro architecture measured with microCT scans at the proximal tibia. Measures are shown as percent change from baseline for trabecular bone volume fraction (Tb.BVTV), trabecular thickness and trabecular separation. Values are means ± SEM; n=13-15/group. Values with different letters are significantly different within time points (Day 14, 28 or 56); lines connect experimental groups (ground control, HLS left, HLS right) that are different from the same experimental group at different time point). All values are significantly different from baseline day zero values (p<0.05).

**Figure 2:**
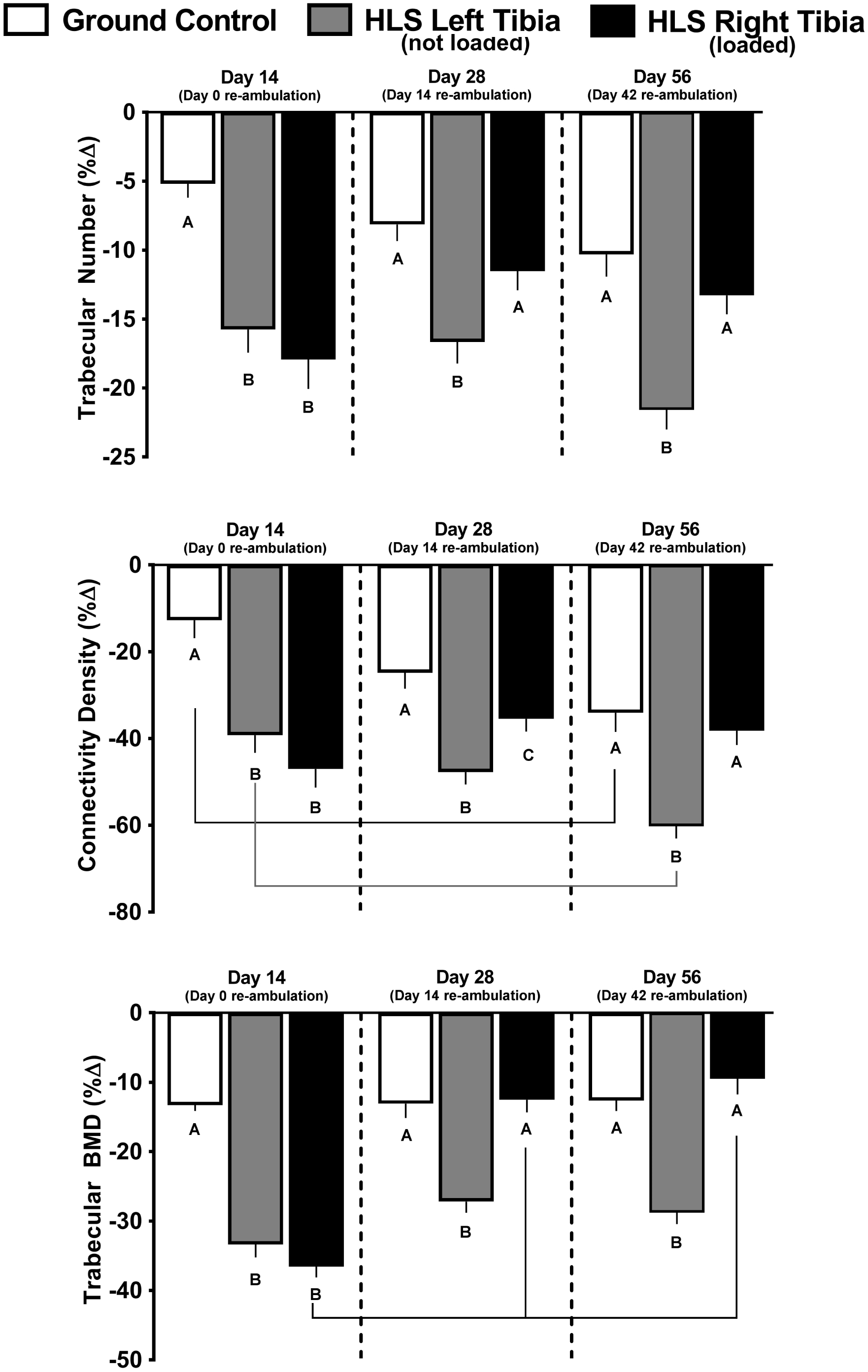
Effect of HLS, re-ambulation only and re-ambulation and mechanical loading on trabecular micro architecture measured with microCT scans at the proximal tibia. Measures are shown as percent change from baseline for trabecular number, trabecular connectivity density and trabecular bone mineral density. Values are means ± SEM; n=13-15/group. Values with different letters are significantly different within time points (Day 14, 28 or 56); lines connect experimental groups (ground control, HLS left, HLS right) that are different from the same experimental group at different time point). All values are significantly different from baseline day zero values (p<0.05).

### Trabecular Bone in HLS Unloaded Mice

Compared to baseline scans at day 0, HLS produced significant bone loss for all trabecular outcomes (**Fig 1, gray bars**). The decrease in trabecular BV/TV, trabecular thickness and increase in trabecular separation are all indicative of catabolic bone changes that were greater than those detected in GC mice from baseline day 0 to day 14 (**Fig 1, gray vs. white bars**). Similarly, the HLS-induced decrease in trabecular number, connectivity density and trabecular bone mineral density was greater than that seen in GC mice (**Fig 2, gray vs. white bars)**.

### Trabecular Bone after Re-ambulation and Acute External Mechanical Loading

During the re-ambulation and loading protocol, there was recovery of some trabecular outcomes as evidence by data from day 28 and 56 (**Fig 1 and 2**). At day 28, in the left limb (re-ambulated but not acutely loaded), trabecular BV/TV showed a partial, non-significant, recovery from day 14 (−37% to -49% from baseline, **Fig 1 gray bar at day 28 vs. gray bar at day 14**), while the right limb (re-ambulated plus acute external mechanical loading) displayed a complete recovery to GC values (−18% to -50% from baseline; **Fig 1 black bar at day 28 vs. black bar at day 14**); these values did not change further at day 56. At day 28, trabecular thickness recovered from day 14 (−11% to -26% from baseline) in the left limb **(Fig 1 gray bar at day 28 vs. gray bar at day 14)**, while the mechanically loaded limb showed a greater recovery (+6 to -28% from baseline (**Fig 1 black bar at day 28 vs. black bar at day 14**). At day 56, the mechanically loaded limb was increased further (+16% to +6% from baseline, **Fig 1 black bar at day 56 vs. black bar at day 28**), while the non-mechanically loaded limb showed no further change **(Fig 1)**. Similarly, acute external mechanical loading in the right limb also reversed the HLS-induced catabolic changes in trabecular separation, trabecular number, trabecular connectivity density and trabecular bone mineral density at days 28 and 56, compared to the contralateral limb that was not exposed to acute external mechanical loading **(Fig 1 & Fig 2)**.

### Cortical Outcomes in Ground Control Mice

Cortical total volume, cortical bone volume, and cortical BV/TV were decreased after 14 days in GC mice, compared to their baseline scans (**Fig 3, white bars**). Furthermore, cortical thickness decreased and cortical bone mineral density increased in GC mice, while cortical porosity was not changed from baseline values (**Fig 4, white bars**). Most GC cortical outcomes did not show further change on day 28 or 56, except for bone mineral density which showed no change from baseline at day 28 (**Fig 4, white bar**).

**Figure 3:**
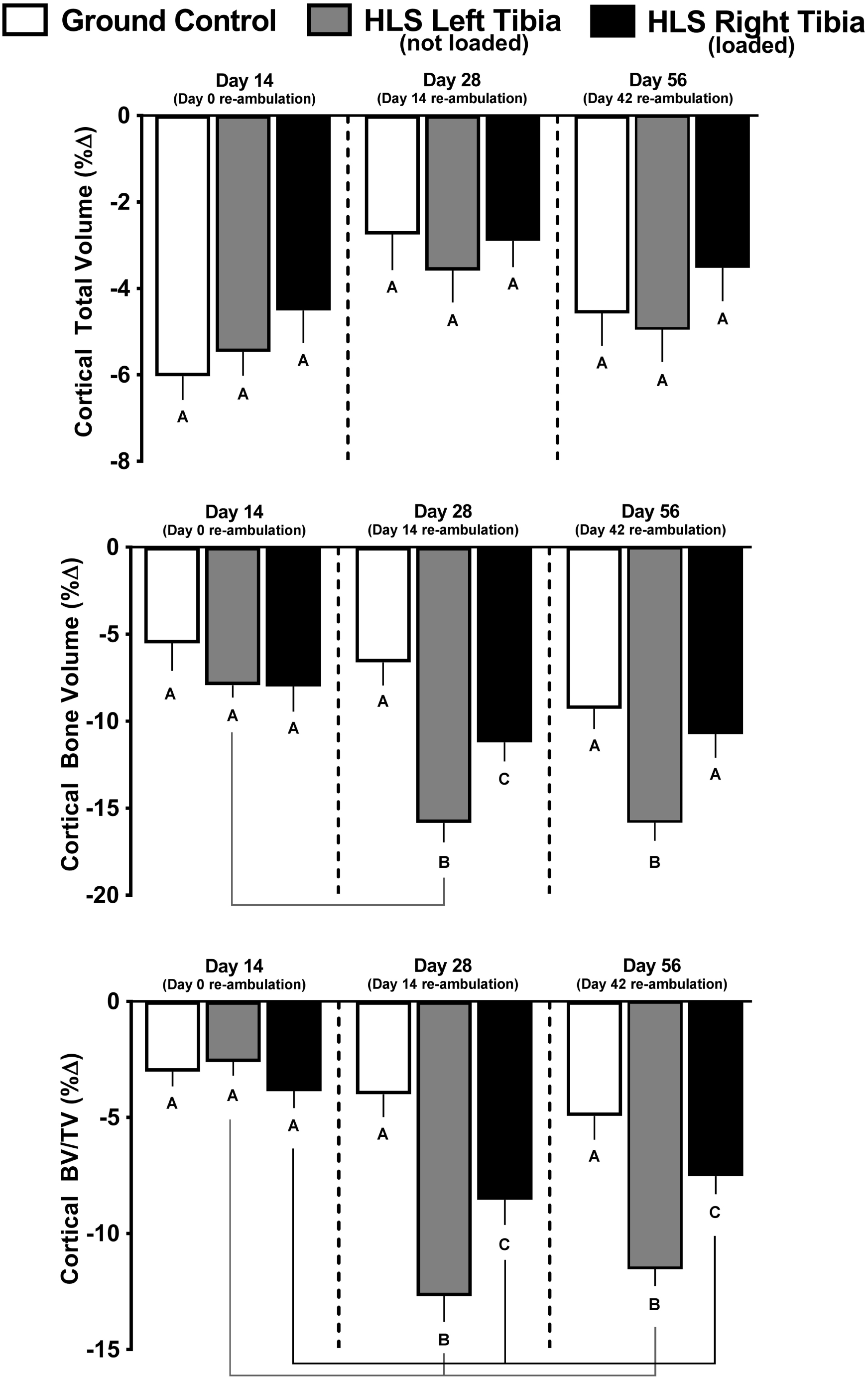
Effect of HLS, re-ambulation only and re-ambulation and mechanical loading on cortical micro architecture measured with microCT scans at the tibia midshaft. Measures are shown as percent change from baseline for cortical total volume, cortical bone volume, and cortical bone volume fraction (BV/TV). Values are means ± SEM; n=13-15/group. Values with different letters are significantly different within time points (Day 14, 28 or 56); lines connect experimental groups (ground control, HLS left, HLS right) that are different from the same experimental group at different time point). All values are significantly different from baseline day zero values (p<0.05).

**Figure 4:**
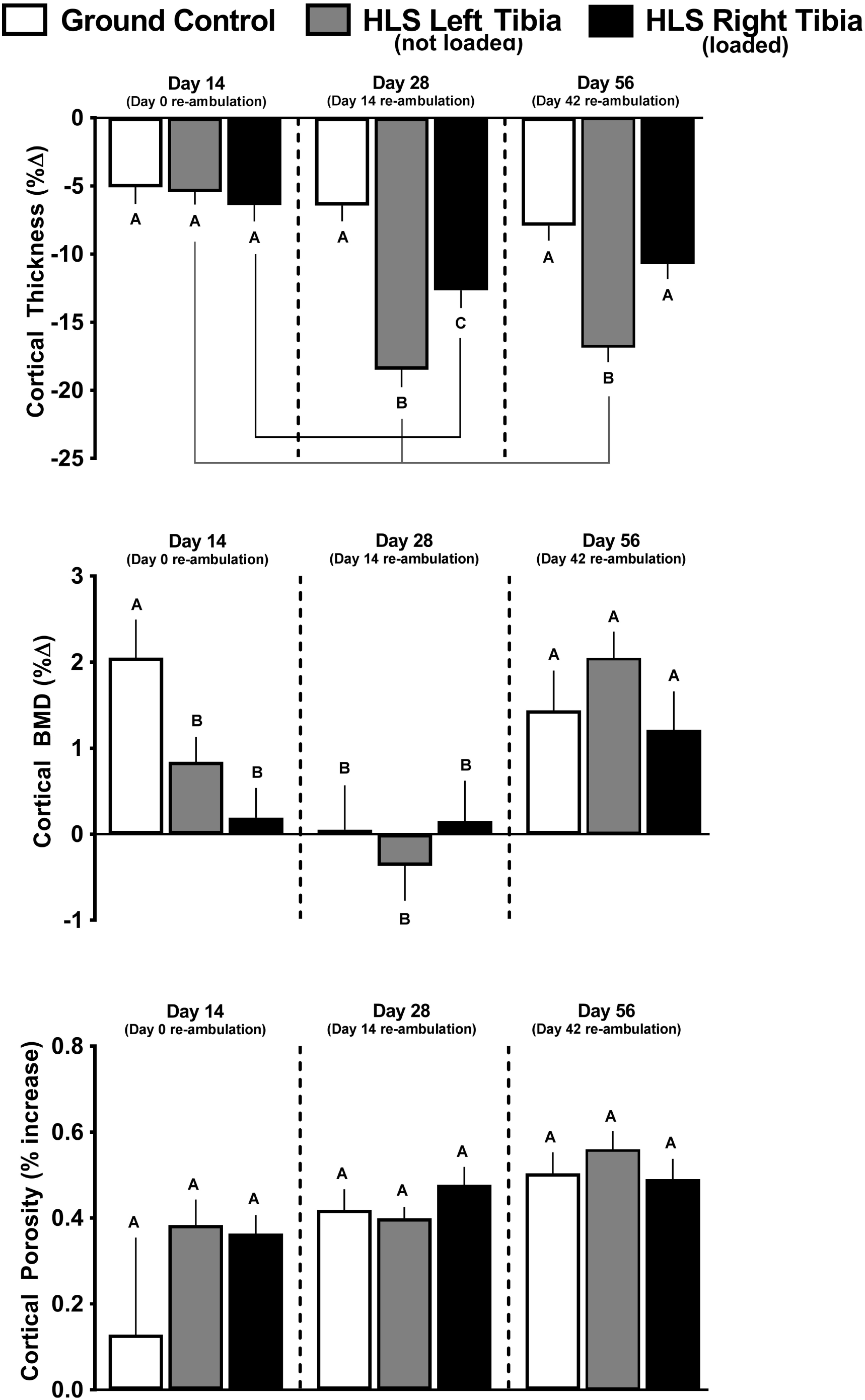
Effect of HLS, re-ambulation only and re-ambulation and mechanical loading on cortical micro architecture measured with microCT scans at the tibia midshaft. Measures are shown as percent change from baseline for cortical thickness, cortical bone mineral density (BMD), and cortical porosity. Values are means ± SEM; n=13-15/group. Values with different letters are significantly different within time points (Day 14, 28 or 56); lines connect experimental groups (ground control, HLS left, HLS right) that are different from the same experimental group at different time point); All values are significantly different from baseline day zero values unless otherwise noted by ♦ symbol (p<0.05).

### Cortical Bone in HLS Unloaded Mice

The HLS-induced decreases in cortical total volume, bone volume and BV/TV were not different from changes seen in GC mice at day 14 **(Fig. 3, gray bars vs. white bars)**. Similarly, HLS-induced decreases in cortical thickness and increases in cortical porosity were similar to GC mice at day 14, while cortical bone mineral density was decreased compared to GC **(Fig 4, gray bars vs white bars)**.

### Cortical Outcomes after Re-ambulation and Acute External Mechanical Loading

There was no effect of re-ambulation (**Fig 3 gray bars**) or acute external mechanical loading (**Fig 3 black bars**) on cortical total volume at either day 28 or 56, relative to GC values at day 28 or 56. Cortical bone volume and cortical BV/TV did exhibit greater catabolic changes at day 28 and 56, compared to GC values (**Fig 3 gray bars vs. white bars**), and this additional loss was mitigated in the limb exposed to acute external mechanical loading compared to the non-mechanically loaded limb (**Fig 3 black bars vs. gray bars**). Similarly, cortical thickness displayed additional loss at day 28 (**Fig 4 gray bars vs. white bars**), and this response was partially attenuated in the limb exposed to acute external mechanical loading (**Fig 4, black bars vs. gray bars**). At day 28 and 56, there was no significant difference in cortical bone mineral density compared to their baseline scans; however, bone mineral density was increased at day 56 compared to baseline (**Fig 4**). Cortical porosity remained increased from baseline values at day 28 and 56, and was not altered by re-ambulation or acute external mechanical loading. (**Fig 4**).

### Muscle Weight and Protein Metabolism

After two weeks, HLS decreased gastrocnemius mass by 15% compared to GC (**Fig 5A, gray bars vs. white bars**). At day 28, the gastrocnemius weight for HLS mice had returned to GC values regardless of whether the limb had been exposed to re-ambulation or re-ambulation in combination with acute external mechanical loading (**Fig 5A, gray and black bars vs. white bars**). At day 56, the left ambulated limb gastrocnemius weight did not differ from GC values, while the right limb was 12% lower (**Fig 5A, gray and black bars vs. white bars**). Results for quadriceps weight showed similar trends to gastrocnemius (data not shown).

**Figure 5:**
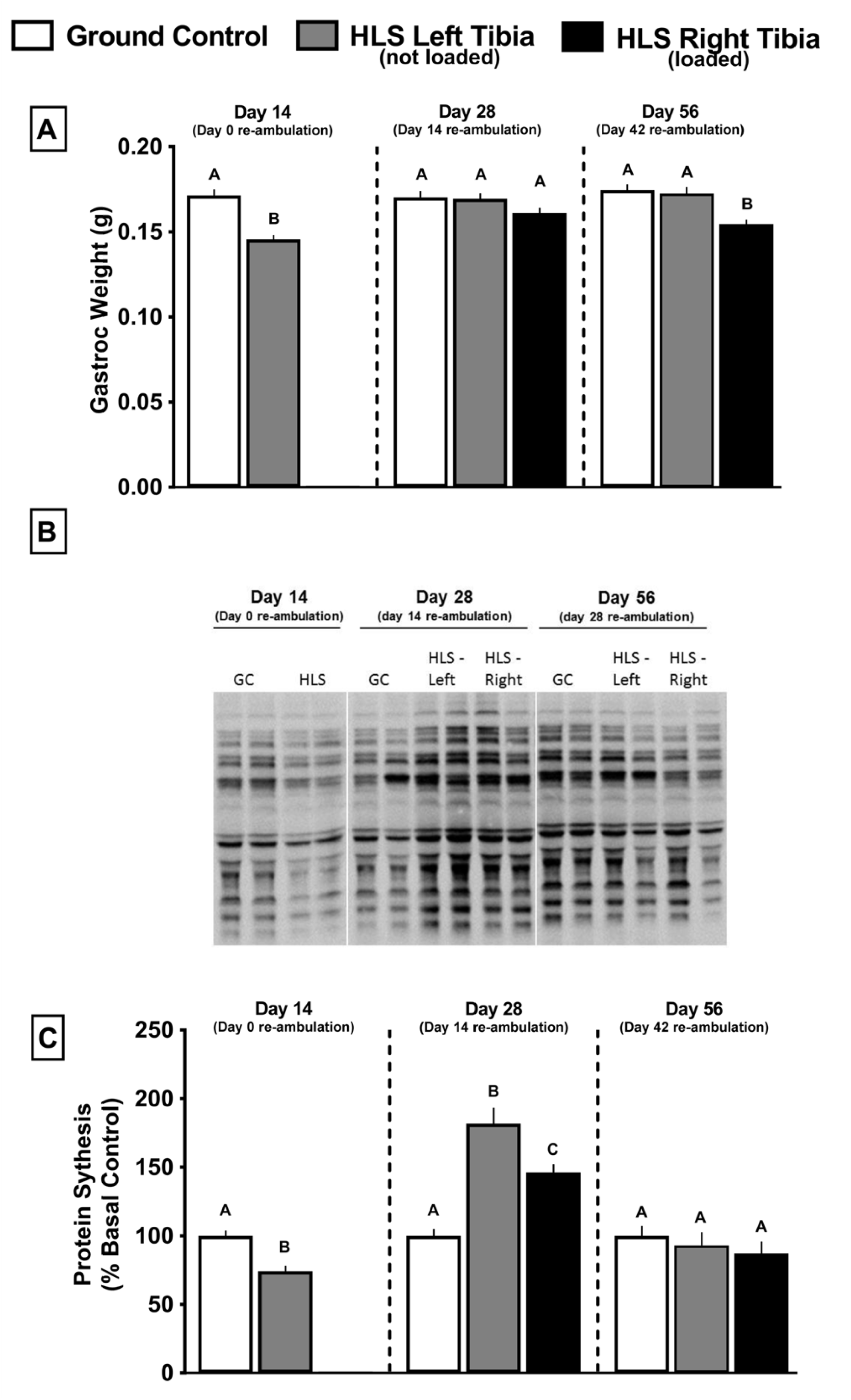
Effect of HLS, re-ambulation only and re-ambulation and mechanical loading on wet muscle weight of gastrocnemius (A), and protein synthesis (C) measured with the SUnSET method. Western blot analysis of the abundance of puromycin-incorporated proteins was determined in muscles from all experimental animals; Panel B is a representative Western blot of puromycin incorporation into muscle protein for different animals in each experimental group at each time point (Day 14, 28 or 56). On day 14, Western analysis was performed only the left muscle from the GC and HLS rats as there was no reason to expect a differential response between the left and right leg at this time point (e.g., no reloading stimulus). Similarly, the GC at day 28 and 56 is also only from the left leg. Vertical white lines depict where the original image was cut to remove gel lanes for presentation purposes. All samples where from a single gel. Densitometry was performed on the entire lane to assess incorporation of puromycin into all protein of various molecular weights. Gels were also stained for Ponceau S and demonstrated equal protein loading (data not shown). Muscle weights are absolute values at time point indicated, and protein synthesis is normalized to GC for each time point. Bar graphs represent means ± SEM; n=13-15/group. Values with different letters are significantly different within time points (p<0.05).

A representative Western blot of puromycin-labeled proteins in the gastrocnemius of rats from each experimental group is illustrated in **Fig 5B**, and the abundance of such proteins was quantitated to provide the relatively rate of in vivo protein synthesis (**Fig 5C**). As a result of HLS, the relative rate of in vivo protein synthesis in the gastrocnemius was decreased 25%, compared to GC mice (**Fig 5C, gray bars vs. white bars**). After 14 days of re-ambulation (day 28), muscle protein synthesis was increased, compared to GC values (**Fig 5C, gray bars vs. white bars**). Ambulation combined with acute external mechanical loading also increased protein synthesis relative to GC, but the increase was smaller than that seen in the re-ambulation alone group (**Fig 5C, black bars vs. gray and white bars**). By day 56, protein synthesis in the gastrocnemius did not differ from the GC mice in either the re-ambulated or re-ambulated with acute external mechanical loading condition (**Fig 5C, black bars vs. gray and white bars**).

After 14 days of unloading, Thr389-phosphorylated S6K1 and Ser240/244-phosphorylated S6 were decreased in the gastrocnemius (**Fig 6B and 6C, gray and black bars vs. white bars**), and these changes were independent of any HLS-induced change in total S6K1 and S6 (**Fig 6A**). In contrast, total and S65-phosphorylated 4E-BP1 did not differ between GC and HLS mice after 14 days of HLS (**Fig 6D, gray bars vs. white bars**). After 14 days of re-ambulation (day 28), phosphorylation of S6K1, S6 and 4E-BP1 were all increased, compared to day 28 GC values (**Fig 6B, C, and D gray bars vs. white bars**). However, at day 28 there was no further increase in phosphorylation of S6K1, S6 or 4E-BP1 in re-ambulated limbs exposed to mechanical loading (**Fig 6B, C, and D black bars vs. gray bars**). At day 56, S6K1 phosphorylation in mice re-ambulated only or re-ambulated and exposed to acute external mechanical loading was similar to that in GC mice (**Fig 6B gray and black bars vs. white bars**). However, at day 56, the phosphorylation of S6 was increased in mice that were re-ambulated alone or re-ambulated in combination with mechanical loading, and the increase was significantly greater in the latter group (**Fig 6C gray and black bars vs. white bars**). Finally, 4E-BP1 phosphorylation on day 56 was increased in re-ambulated only mice, but not in re-ambulated mice exposed to acute external mechanical loading, relative to GC mice (**Fig 6 gray and black bars vs. white bars**).

**Figure 6:**
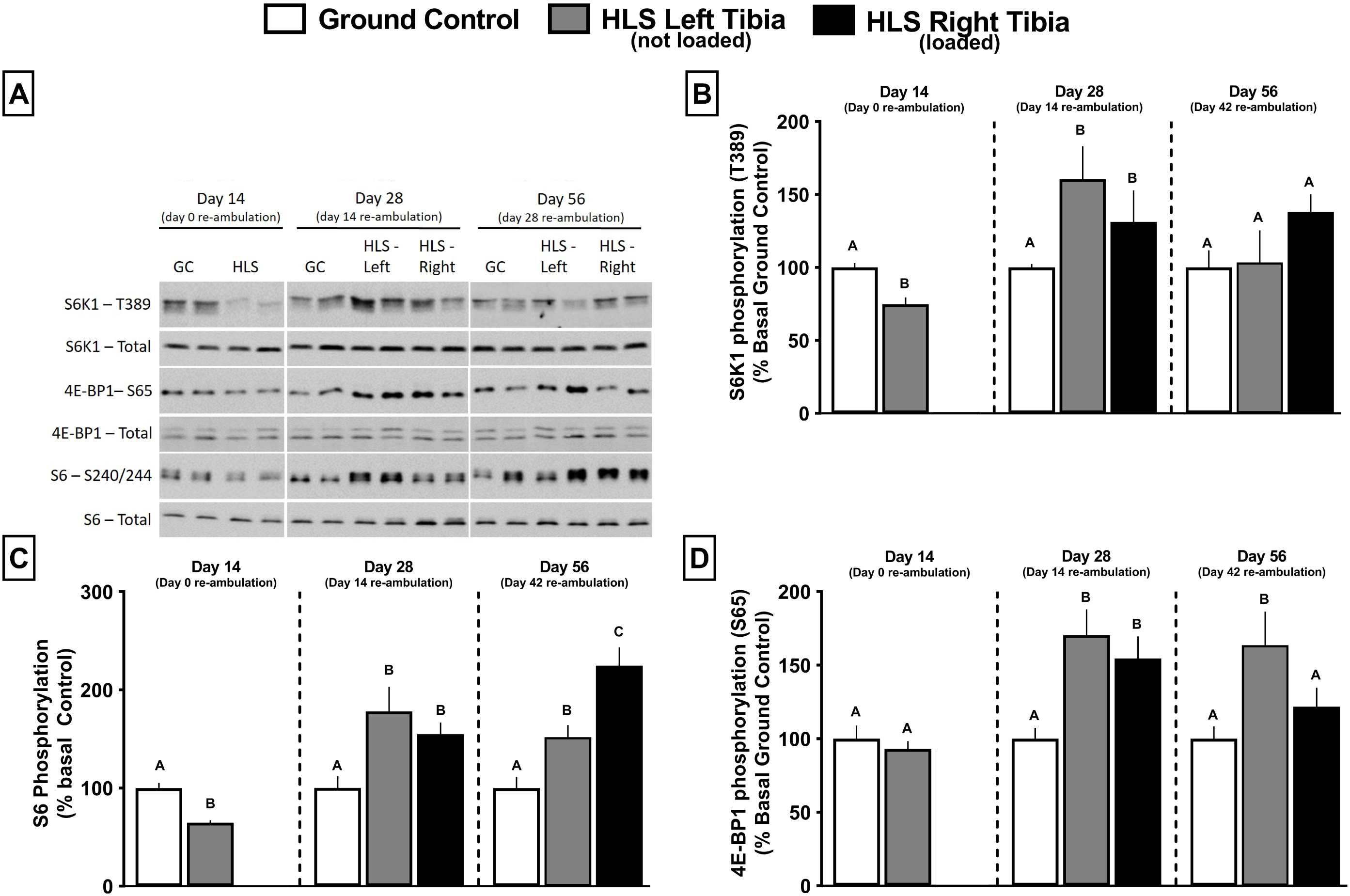
Effect of HLS, re-ambulation only and re-ambulation and mechanical loading on the phosphorylation of mTOR downstream regulators 4EBP1 and S6K1 measured via Western blot analysis. Western blot analysis was performed on muscles from all rats and the top left panel (A) is a representative composite image for the relative abundance of each total and phosphorylated protein assessed for each experimental group at each time point (Day 14, 28 or 56). On day 14, Western analysis was performed on only the left muscle from the GC and HLS rats as there was no reason to expect a differential response between the left and right leg at this time point (e.g., no reloading stimulus). Similarly, the GC at day 28 and 56 is also only from the left leg. Vertical white lines depict where original images were cut to remove gel lanes for presentation purposes. Western blots for the same protein were all run on the same day and processed similarly. Samples for each experimental day are from a single gel. Panels B, C and D show phosphorylation of S6K1, S6 and 4E-BP1, respectively, in gastrocnemius from each experimental group at each time point (Day 14, 28 or 56); all samples for a given time point were run on the same gel. Bar graphs represent average values at conclusion time points, and are first normalized the amount of the respective nonphosphorylated protein and then normalized to the GC group for each time point. Bar graphs represent means ± SEM; n=13-15/group. Values with different letters are significantly different within time points (p<0.05).

## Discussion

Our results suggest that acute external mechanical loading, of a duration and load previously shown to induce site-specific increases in mineral content [26], facilitates bone recovery following a period of HLS. At day 14, there was detectable trabecular bone loss in GC mice, due in part to age, but with substantial additional loss observed in HLS mice. The age-related decreased trabecular BV/TV (approximately 20%) in GC mice from 15 weeks (start of the experiment) to 17 weeks was similar to the 30% decrease from 12 to 15 weeks of age reported by Glatt et al [21], the approximately 10% decrease reported by Halloran et al. [22], and the approximately 15% decrease reported by Lloyd et al [23]. Despite this additional unloading-induced bone loss, all of the trabecular outcomes in the limb of HLS mice demonstrated a full or partial recovery to GC values by day 28 (2 weeks of re-ambulation and loading). This included a full recovery of BV/TV. Furthermore, there was recovery to GC values for trabecular separation, number, and bone mineral density, while trabecular connectivity density showed a partial recovery compared to the contralateral limb that did not receive acute external mechanical loading. For most of these outcomes, additional effects of loading were not observed at day 56, suggesting that trabecular bone adaptation in mice occurs mostly within 2 weeks after the start of acute external mechanical loading. Our data demonstrating acute external mechanical loading-induced recovery of trabecular bone lost to HLS, within two weeks, suggests that trabecular bone sensitivity to loading, and the subsequent anabolic response, is similar to the sensitivity to unloading, and subsequent catabolic response. These data suggest that acute loading has the potential to fully mitigate the catabolic effects of unloading, and that unloading does not affect the ability of bone to respond to subsequent acute external mechanical loading.

In contrast to trabecular data, two weeks of HLS did not increase the aging-induced loss of cortical bone in suspended mice, compared to loss in their GC counterparts. However, several cortical parameters (including cortical bone volume, cortical BV/TV, and cortical thickness) did show catabolic changes after HLS, although not until 14 days of re-ambulation. Furthermore, for all three of these parameters, acute external mechanical loading during re-ambulation mitigated HLS-induced bone loss. These data suggest a delayed response of cortical bone to unloading that has not been previously reported. Our findings confirm previous studies from our group demonstrating that two weeks of HLS does not alter cortical outcomes compared to GC measured immediately following unloading [24]. However, in other studies we have demonstrated that three weeks of HLS does result in additional cortical bone loss [23, 25], indicating that cortical bone is less responsive to unloading than trabecular bone.

The differential sensitivity of cortical and trabecular bone to the catabolic effects of HLS appear to be due, at least in part, to the slower turnover of cortical bone. This supposition is supported by the observation that cortical bone in BL/6 mice continues to grow until at least three months of age and thereafter does not change with age [21, 22], which could mask the effects of unloading. In contrast, trabecular bone loss in BL/6 mice begins at 6-8 weeks of age and continues thereafter [21, 22].

The reduction in muscle protein synthesis after two weeks of HLS is consistent with previous reports from our laboratory [23]. Decreases in protein synthesis are concomitant with decreases in muscle mass at the same time point (day 14). This decrease in protein synthesis appears at least partially due to reduced mTORC1 kinase activity as evidenced by the reduction in its downstream substrate, S6K1. Furthermore, the HLS-induced decrease in S6K1 activity is consistent with the reduced phosphorylation of one of its substrates, the ribosomal protein S6. In contrast, at day 14, the mTORC1 downstream substrate 4EBP1 was not altered, a result that is consistent with previous data from our group showing S6K1 changes precede 4EBP1 changes following HLS [23].

The increase in muscle mass following re-ambulation was accompanied by an increase in protein synthesis that was present after 14 days of re-ambulation (day 28). However, despite the smaller increase in protein synthesis in the re-ambulated limbs and limbs exposed to acute external mechanical load, compared to the re-ambulated only limb, muscle mass did not differ. Collectively, these data suggest that the rate of protein degradation, either through the ubiquitin-proteasome pathway or autophagy, is lower in re-ambulated limbs and limbs exposed to acute external mechanical load such that a smaller increment in protein synthesis leads to the same increase in muscle mass. Regardless of the potential contribution of protein degradation, this decreased rate of protein synthesis in the right limb with external loading compared to the left limb without loading could not be explained by a difference in the phosphorylation of S6K1/S6 or 4E-BP1 suggesting that mTOC1 activity did not differ at day 28. The underlying mechanism for this diminished rate of protein synthesis in response to external mechanical loading was not assessed and is unknown, but conceptually could result from the differential activation of satellite cells.

At day 56, consistent with the decreased protein synthesis observed at day 28, gastrocnemius mass was decreased in the re-ambulated and loaded right limb relative to the re-ambulated only left limb, reinforcing the concept that changes in protein synthesis precede changes in muscle mass. To that end, protein synthesis returned to levels seen in GC mice at day 56 in both limbs. Despite this unaltered rate of synthesis, increased phosphorylation of S6 and 4E-BP1 were still detected at this latter time point. The reason for this apparent mismatch between synthesis and mTORC1 activity is unknown.

This study examined changes in bone and muscle following unloading, unloading and re-ambulation, and unloading and re-ambulation combined with acute external mechanical loading. One limitation of this study is that it only examined male mice. However, previous studies suggest a similar anabolic response of male and female C57Bl/6 mice to tibial loading [26]. Thus, including female mice in this study would likely not have changed our interpretation of the results. Our results suggest that muscle responds to acute loading following re-ambulation differently than bone. While re-ambulation alone attenuated both muscle and bone loss resulting from HLS, acute external mechanical loading further attenuated bone, but not muscle loss following HLS and re-ambulation. Additionally, by day 56 there was a negative effect of acute external mechanical loading on muscle mass. Thus, our data do not support the concept that acute external mechanical loading facilitates recovery of bone and muscle loss resulting from unloading and suggest that bone and muscle do not respond to their mechanical environment as a fully integrated unit.

## Materials and Methods

### Animal Procedures

Male C57BL/6J mice (Jackson Laboratories, Bar Harbor, ME, USA) were used for all experiments. All mice used were approximately 105 days (15 weeks) old ± 3 days and thus skeletally mature on day 0 (start of experiment). Mice were housed in standard polycarbonate enclosures modified for HLS (2 mice/cage), with room temperature at 25°C and on a 12 hour light/dark cycle. Standard rodent 2018 Tekland Global 18% protein rodent diet (Harlan Laboratories Inc., Indianapolis, IN, USA) and water were provided ad libitum throughout the study. Mice were acclimated to the room used for HLS and allowed to ambulate normally for 2 weeks prior to study commencement. Mice were randomly assigned to the HLS group, HLS and acute externally loaded group, or the time-matched ground control (GC) group, with 13-15 mice in each group. All procedures followed NIH guidelines for experimental animals, and were approved by the Penn State College of Medicine Institutional Animal Care and Use Committee (Protocol # 46451).

### Hind Limb Suspension Mechanical Unloading

We used a modified model of HLS originally described by Morey-Holton and Globus [27], and previously reported by our lab [23-25]. Mice were anesthetized with isoflurane (2% + oxygen) and a baseline microCT scan was performed prior to HLS. To suspend the mice, two tape strips were applied to the tail that secured a string that was attached to a metal bar across the top of the cage. Strings were adjusted to support the mouse at 30 degrees of elevation, which is adequate for consistent HLS unloading while avoiding unnecessary strain on the animal. Mice were suspended in cages as pairs; however, tethering to the suspension apparatus prevented physical contact. Following 2 weeks of unloading, HLS mice were returned to re-ambulation in cages identical to those of GC mice (one mouse/cage). GC mice were placed alone (one mouse/cage) in standard mouse cages for the duration of the study and were not suspended. HLS mice received urethral cleaning twice per day using saline and alcohol wipes to prevent formation of plugs, a common observation in HLS male mice.

### Tibia Compression Mechanical Loading

Non-invasive compressive loading of the right tibia began immediately following the 2-week HLS protocol, and continued for 6 weeks at 4 consecutive days/week [28]. The right leg of anesthetized (2% isoflurane + oxygen) and supine mice were positioned horizontally in line with a plastic cup covered with 1/8 inch foam padding. The cup encapsulated the flexed knee, while a rigid plastic fixture held the foot at 30 degrees dorsiflexion, as reported by Fritton, et al [29]. Each tibia was loaded for 1,200 cycles/day with 9N compression in a saw-tooth waveform at 4 Hz, including 0.1 s dwell at 2.0 N between cycles. Therefore, the total length of each loading protocol was approximately 5 minutes. Previous studies examining strain at tibial mid-shaft in C57Bl/6J mice reported a linear relationship between load and microstrain (μϵ) using loads ranging from 3.8-11.3N [30]. Therefore, our load of 9N corresponded to tibial deformations of about 1400 μϵ at our site of mid-shaft scanning. While this load is less than typically utilized in compressive loading studies [28], those other studies did not mechanically load bone that had previously been subjected to the catabolic state produced by HLS. The left leg of HLS mice was used as an internal control and did not receive mechanical compression. Time-matched control mice not receiving loading were anesthetized (2% isoflurane + oxygen) for 5 mins on the daily loading schedule. Acute external mechanical loading did not appear to affect gait and it did not appear that the mice favored the loaded limb.

### Micro-Computed Tomography

Right and left tibias were scanned using a Scanco vivaCT 40 microCT (Scanco Medical AG, Bruttsellen, Switzerland) on day 0 as a baseline, and days 14, 28, or 56. Baseline scans were performed in vivo, and subsequent scans were performed on bones that had been wrapped in phosphate buffered saline (PBS) soaked gauze, placed in 1.5 ml tubes and frozen at -80°C following removal from the animal. Trabecular sections from proximal tibia were evaluated over a 72 slice region, as previously reported [23-25]. Cortical sections from the mid-shaft of the tibia were evaluated over a 22-slice region. Settings were 55 KVp, 145 µa, and 200 ms integration time. Image reconstruction was 2048 × 2048 x 76 isotropic voxels 10.5 µm wide. Images were Gaussian filtered (sigma=1.5, support=2) and a 27.5% threshold was used to remove soft tissue. 72 slice trabecular regions were manually segmented with automated contouring between manually marked regions of interest. Cortical 22 slice region outlines (periosteal and endosteal boundaries) were segmented with a semi-automated edge detecting sequence using Scanco evaluation software. Data were analyzed in a blinded manner according to previous guidelines (40). Trabecular parameters included bone volume percentage (Tb. BV/TV), number (Tb.N), thickness (Tb.Th), separation (Tb.Sp), connectivity density (Conn. D), and bone mineral density (TBMD). Cortical parameters included total volume (TV), total bone volume (BV), bone volume fraction percentage (Ct. BV/TV), thickness (Ct. Th), porosity (Ct. Po), and bone mineral density (Ct. BMD).

### Protein Synthesis

In vivo muscle protein synthesis was assessed using the non-isotopic SUnSET method and an antibody against puromycin (Kerafast, Boston, MA), as described previously [31, 32]. All animals received an intraperitoneal injection of 0.04 μmol/g body weight of puromycin dissolved in sterile saline 30 min prior to tissue collection. Muscles (gastrocnemius and quadriceps) were subsequently excised, frozen with liquid nitrogen-cooled clamps, weighed, and stored at -80°C. Western blotting was performed for the immunological detection of puromycin-labeled peptides that are synthesized during the 30-min period between injection and euthanasia [31, 32]. As reviewed, the SUnSET method is a valid and accurate alternative to the use of radioisotopes for determining protein synthesis in vitro and in vivo in muscle, although this method is limited in being able to only determine relative (not absolute) rates of protein synthesis [31, 32].

### Western Blot Analysis

The phosphorylation state of 4E-BP1 and S6K1 has been typically used to assess the in vivo activation of mTORC1 (mechanistic target of rapamycin complex 1) which is a protein kinase that is central in regulating protein synthesis via various different inputs (e.g., growth factors, hormones, energy status, and mechanical stress) [33]. Thus, changes in the phosphorylation of 4E-BP1 and S6K1 as well as S6 (which is phosphorylated by S6K1) are directly proportional to changes in muscle protein synthesis. To assess phosphorylation of 4E-BP1, S6K1 and S6, muscle was homogenized in ice-cold buffer, protein concentration quantified and equal amounts of protein subjected to SDS-PAGE. Western analysis was performed in a blinded manner for Ser65-phosphorylated 4E-BP1 (#9451; Cell Signaling Technology (CST), Boston, MA), total 4E-BP1, (gift of Scot Kimball, Penn State College Medicine), Thr389-phosphorylated S6K1 (#9205; CST), and total S6K1 (#SC-230; Santa Cruz Biotechnologies, Dallas, TX), Ser240/244-phosphorylated S6 (#2215; CST), and total S6 (#2217; CST), and puromycin (described above). Blots were developed with enhanced chemiluminescence reagents (Supersignal Pico, Pierce Chemical; Rockford, IL). Dried blots were exposed to x-ray film to achieve a signal within the linear range, scanned, and quantified using Scion Image 3b software (Scion Corp; Frederick, MD). Samples from all six experimental groups were run on the same gel and phosphorylated levels are expressed relative to total amounts of the respective protein. The sample size for each group and time was 13-15, and these samples were run on 3 independent occasions.

### Statistical analysis

Data are shown as mean ± standard error. Data were analyzed with GraphPad Prism (La Jolla, CA) using a one-way ANOVA with Student-Newman-Keuls post hoc analysis when 3 groups where compared. Statistical significance was set at p<0.05.

## Data Availability

All data generated or analyzed during this study are included in this published article or if not are available from the corresponding author on reasonable request

## Acknowledgements

This work was supported by National Space Biomedical Research Institute grant MA02802, and NIH grant R01AG13097.

## Competing interests

The authors declare no potential conflicts of interest, financial or otherwise.

## Author Contributions

**AR Krause:** Designed experiments, performed hind limb unloading, performed acute mechanical loading, completed bone analyses, statistical analyses of data, interpreted data, and wrote initial drafts of manuscript.

**TA Speacht:** Performed hind limb unloading and completed bone analyses.

**JL Steiner:** Completed muscle analyses, statistical analyses, and wrote parts of manuscript.

**CH Lang:** Completed muscle analyses, statistical analyses, interpreted data, wrote parts of manuscript, and edited final drafts.

**HJ Donahue:** Designed experiments, interpreted data, wrote final manuscript draft.

## Notes

### Competing Interest Statement

The authors have declared no competing interest.

