## Supplemental Fig 1 for "Mechanical Loading Recovers Bone but not Muscle Lost During Unloading"

### Slide 1
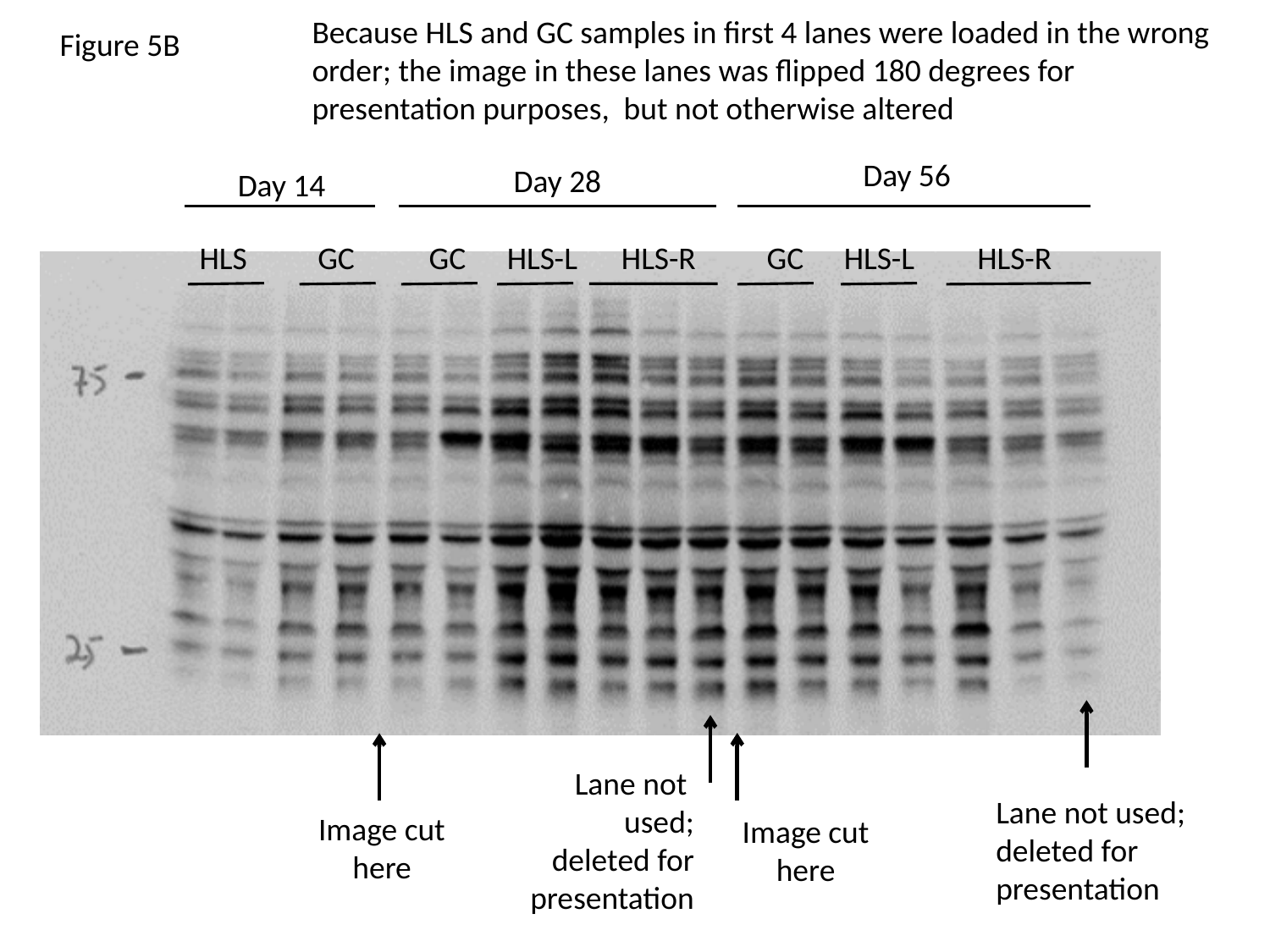

Because HLS and GC samples in first 4 lanes were loaded in the wrong order; the image in these lanes was flipped 180 degrees for presentation purposes, but not otherwise altered
Figure 5B
Day 56
Day 28
Day 14
HLS
GC
GC
HLS-L
HLS-R
GC
HLS-L
HLS-R
Lane not
used;
deleted for
presentation
Lane not used;
deleted for
presentation
Image cut
here
Image cut
here

### Slide 2
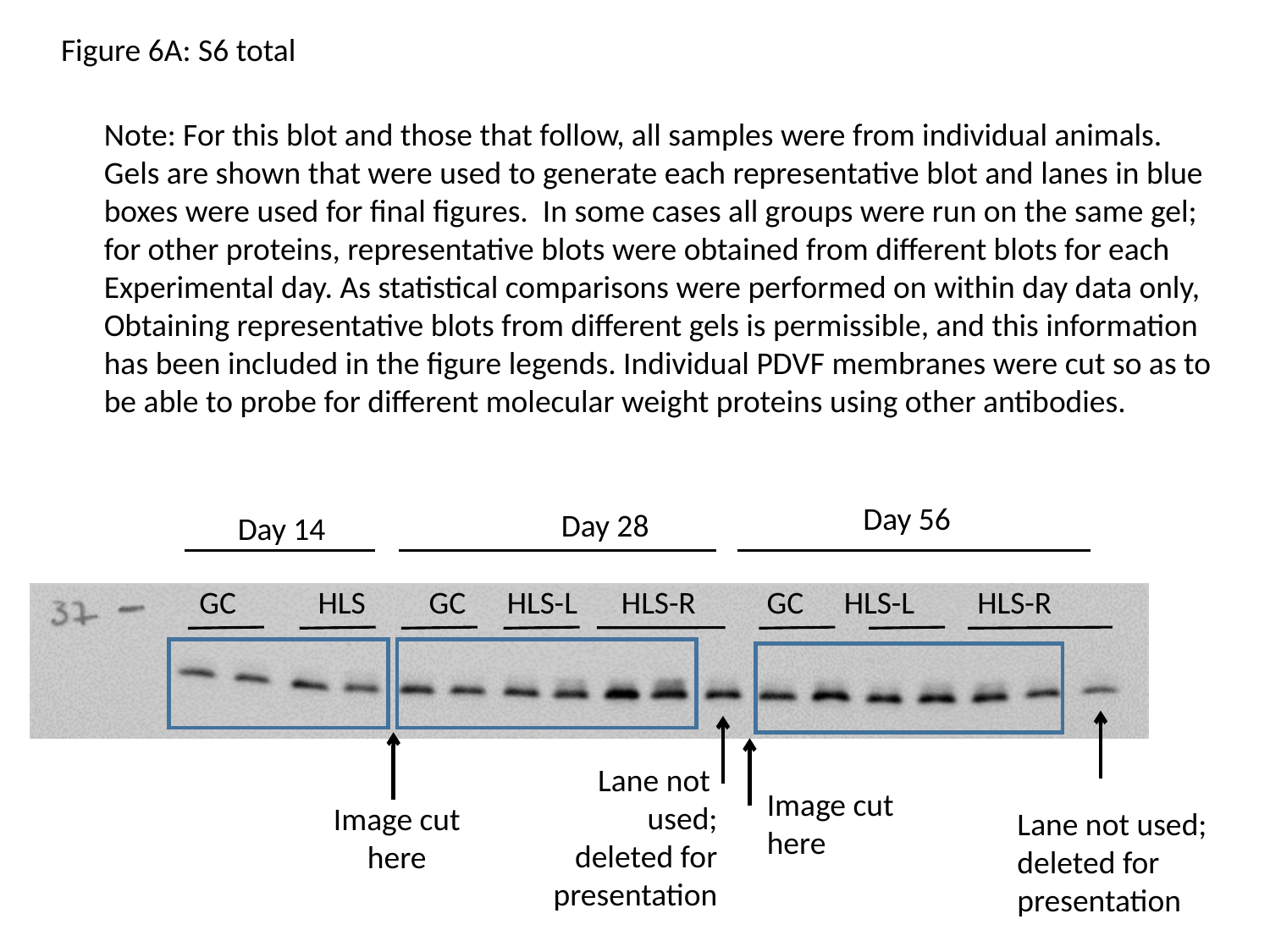

Figure 6A: S6 total
Note: For this blot and those that follow, all samples were from individual animals.
Gels are shown that were used to generate each representative blot and lanes in blue
boxes were used for final figures. In some cases all groups were run on the same gel;
for other proteins, representative blots were obtained from different blots for each
Experimental day. As statistical comparisons were performed on within day data only,
Obtaining representative blots from different gels is permissible, and this information
has been included in the figure legends. Individual PDVF membranes were cut so as to
be able to probe for different molecular weight proteins using other antibodies.
Day 56
Day 28
Day 14
GC
HLS
GC
HLS-L
HLS-R
GC
HLS-L
HLS-R
Lane not
used;
deleted for
presentation
Image cut
here
Image cut
here
Lane not used;
deleted for
presentation

### Slide 3
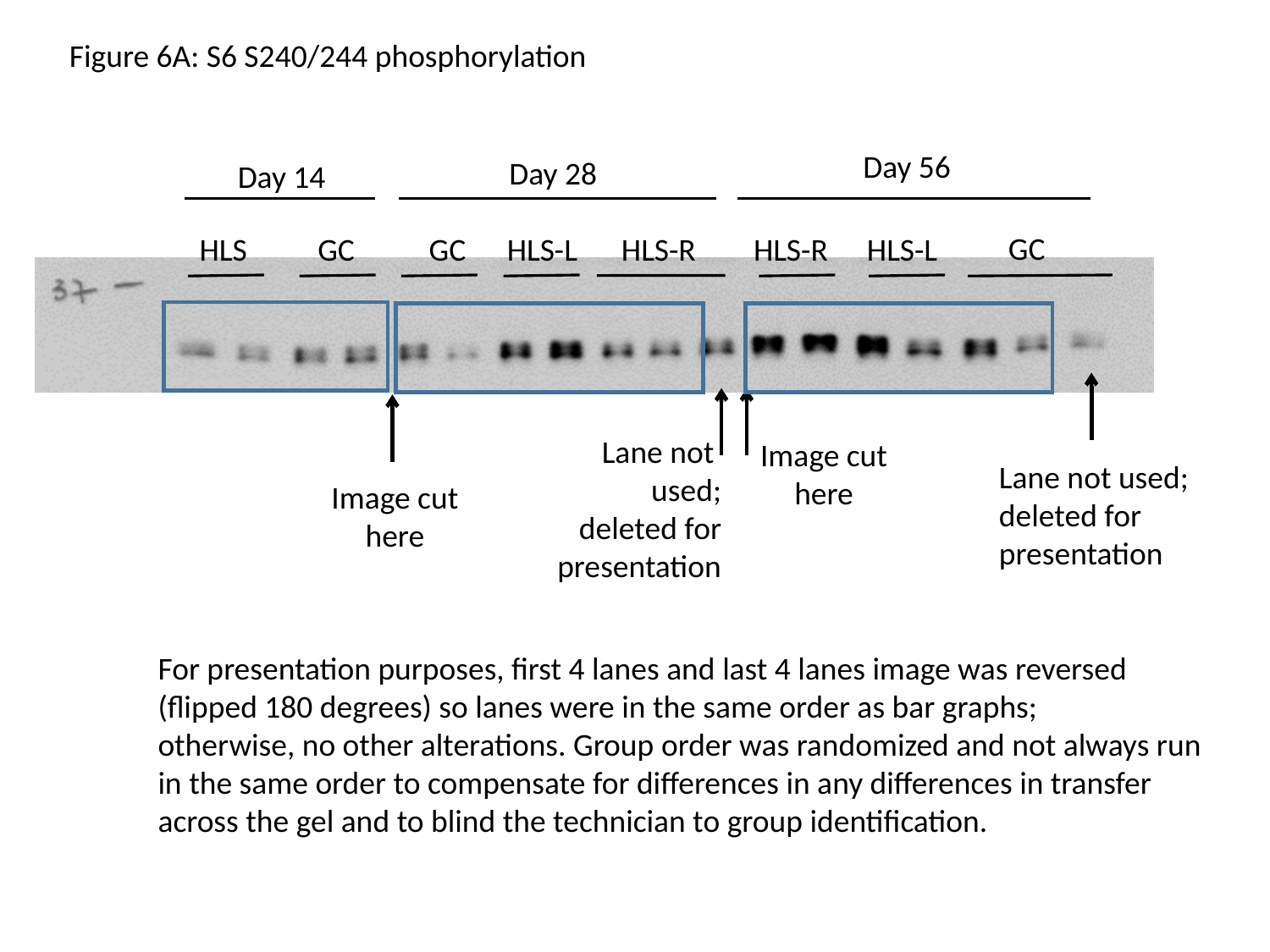

Figure 6A: S6 S240/244 phosphorylation
Day 56
Day 28
Day 14
GC
HLS
GC
GC
HLS-L
HLS-R
HLS-R
HLS-L
Lane not
used;
deleted for
presentation
Image cut
here
Lane not used;
deleted for
presentation
Image cut
here
For presentation purposes, first 4 lanes and last 4 lanes image was reversed
(flipped 180 degrees) so lanes were in the same order as bar graphs;
otherwise, no other alterations. Group order was randomized and not always run
in the same order to compensate for differences in any differences in transfer
across the gel and to blind the technician to group identification.

### Slide 4
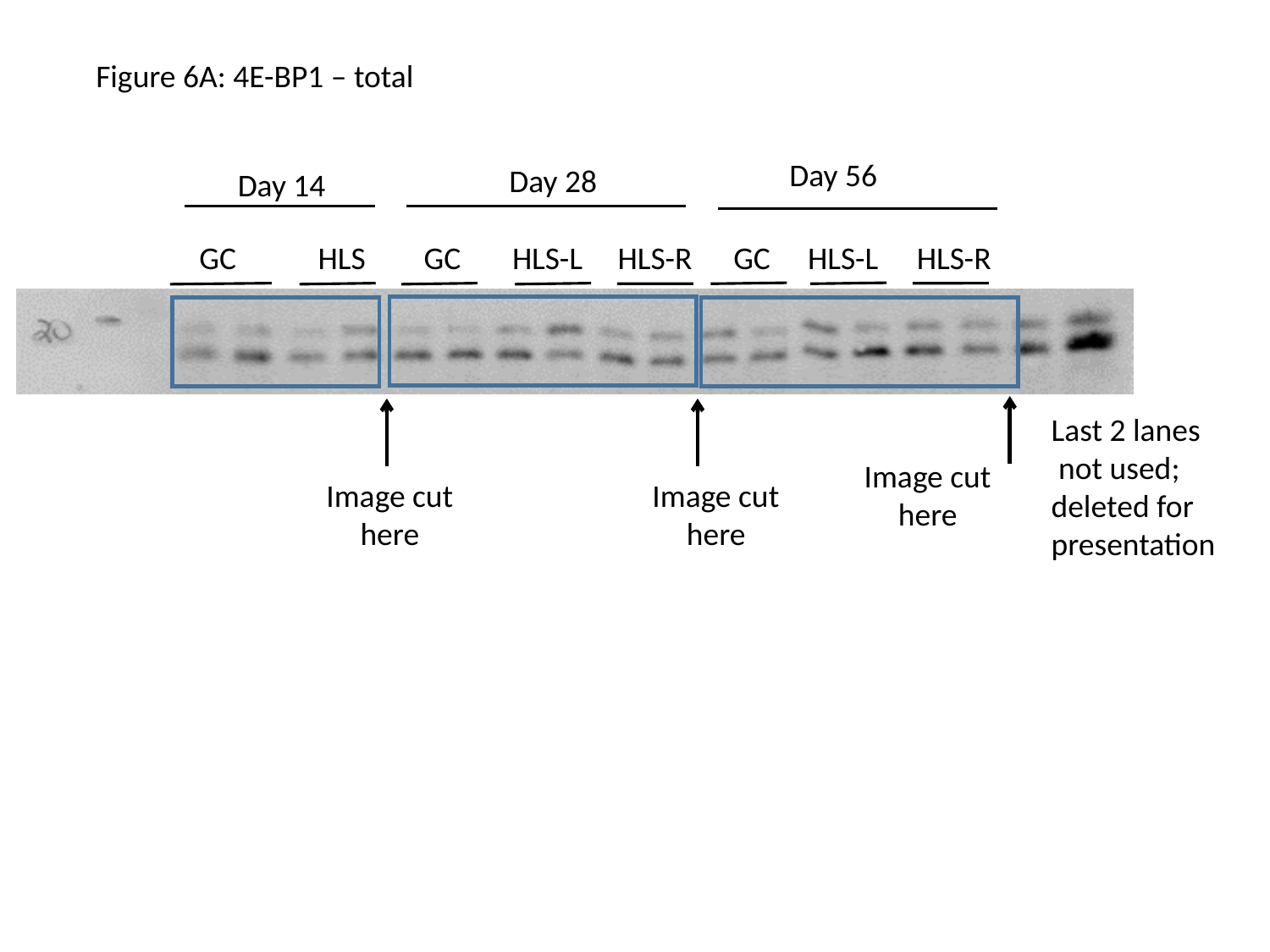

Figure 6A: 4E-BP1 – total
Day 56
Day 28
Day 14
GC
HLS-L
HLS-R
GC
HLS
GC
HLS-L
HLS-R
Last 2 lanes
 not used;
deleted for
presentation
Image cut
here
Image cut
here
Image cut
here

### Slide 5
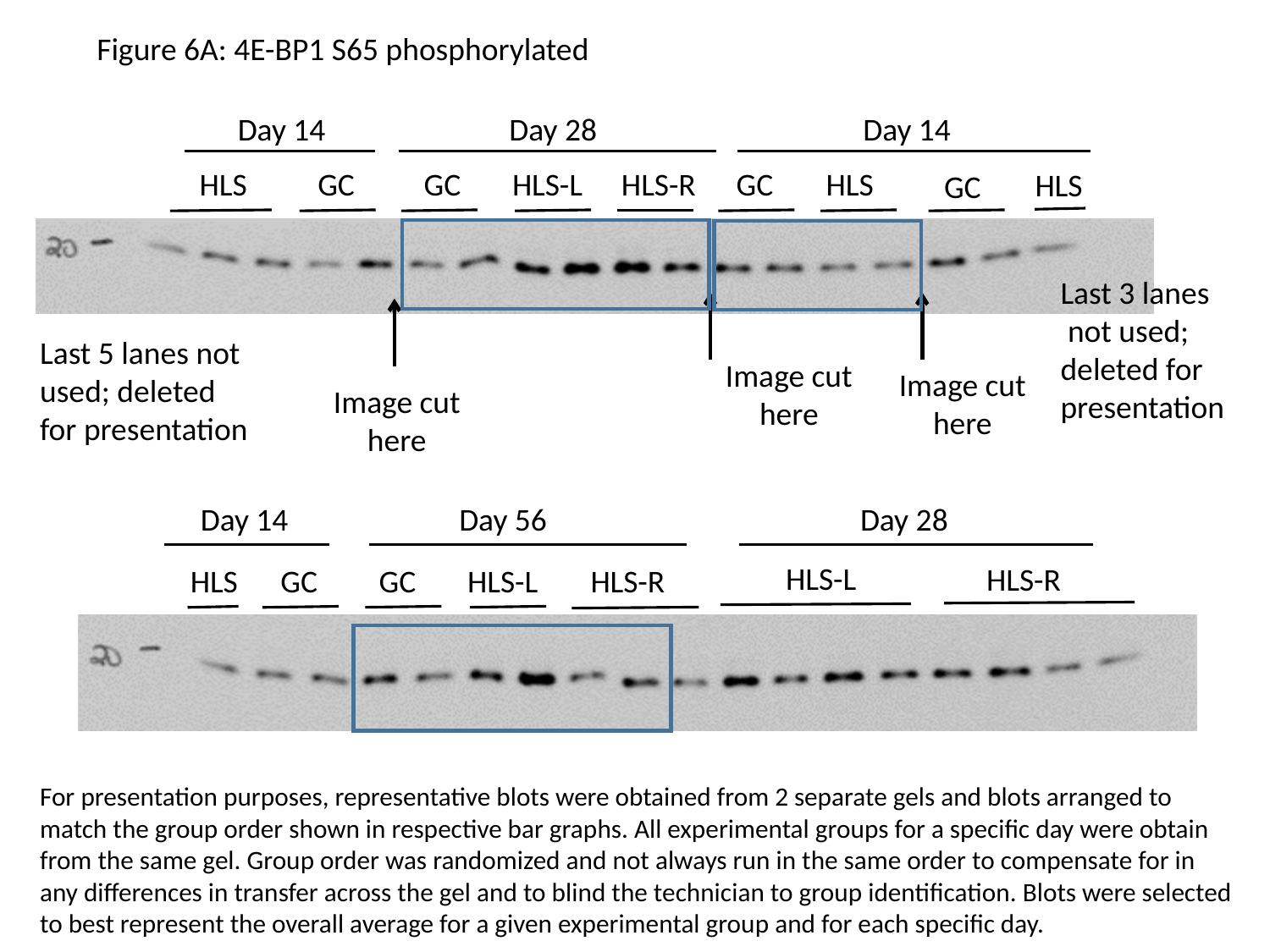

Figure 6A: 4E-BP1 S65 phosphorylated
Day 14
Day 28
Day 14
HLS
GC
GC
HLS-L
HLS-R
GC
HLS
HLS
GC
Last 3 lanes
 not used;
deleted for
presentation
Last 5 lanes not used; deleted for presentation
Image cut
here
Image cut
here
Image cut
here
Day 14
Day 56
Day 28
HLS-L
HLS-R
GC
HLS-L
HLS-R
HLS
GC
For presentation purposes, representative blots were obtained from 2 separate gels and blots arranged to match the group order shown in respective bar graphs. All experimental groups for a specific day were obtain from the same gel. Group order was randomized and not always run in the same order to compensate for in any differences in transfer across the gel and to blind the technician to group identification. Blots were selected to best represent the overall average for a given experimental group and for each specific day.

### Slide 6
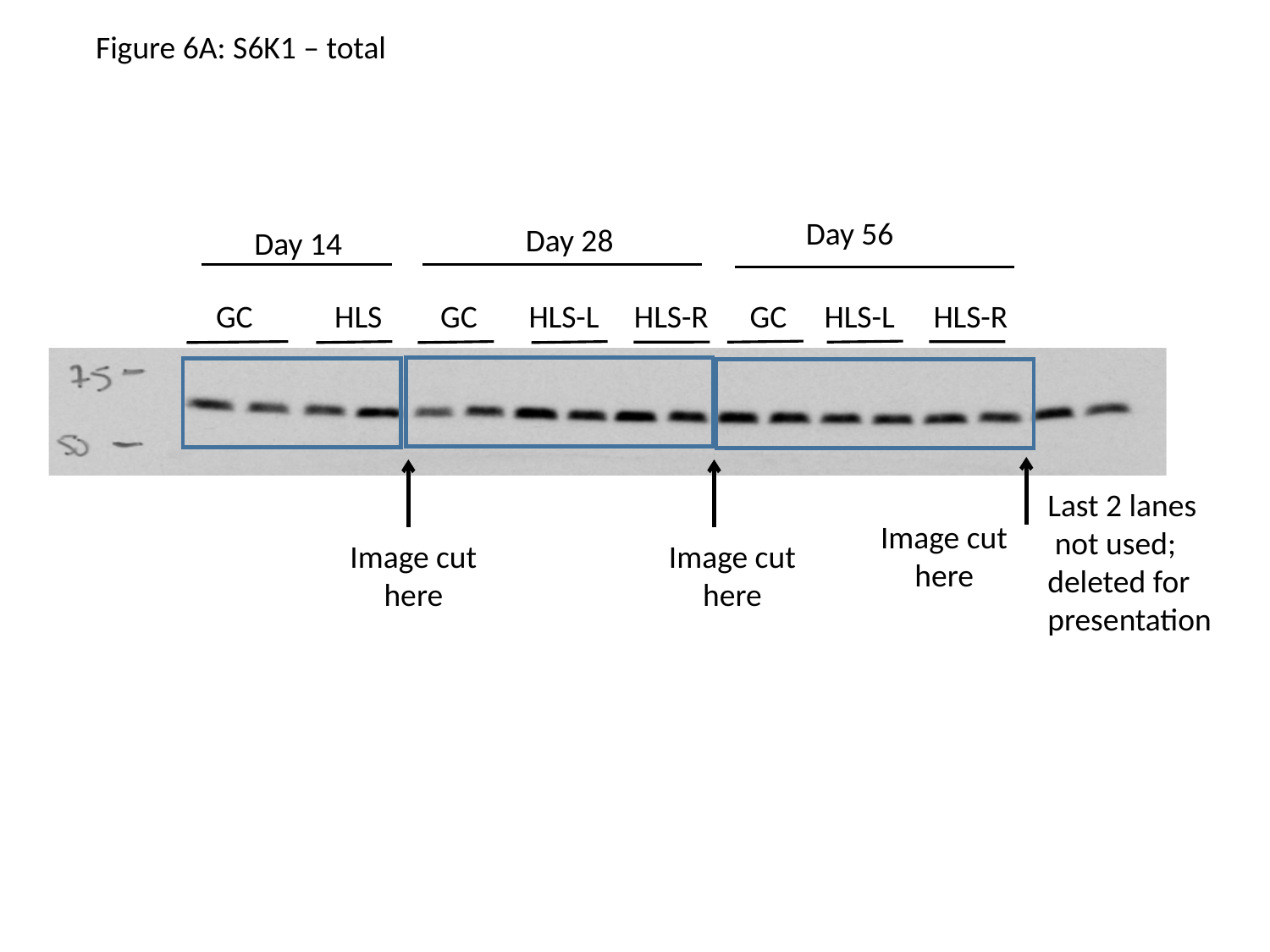

Figure 6A: S6K1 – total
Day 56
Day 28
Day 14
GC
HLS-L
HLS-R
GC
HLS
GC
HLS-L
HLS-R
Last 2 lanes
 not used;
deleted for
presentation
Image cut
here
Image cut
here
Image cut
here

### Slide 7
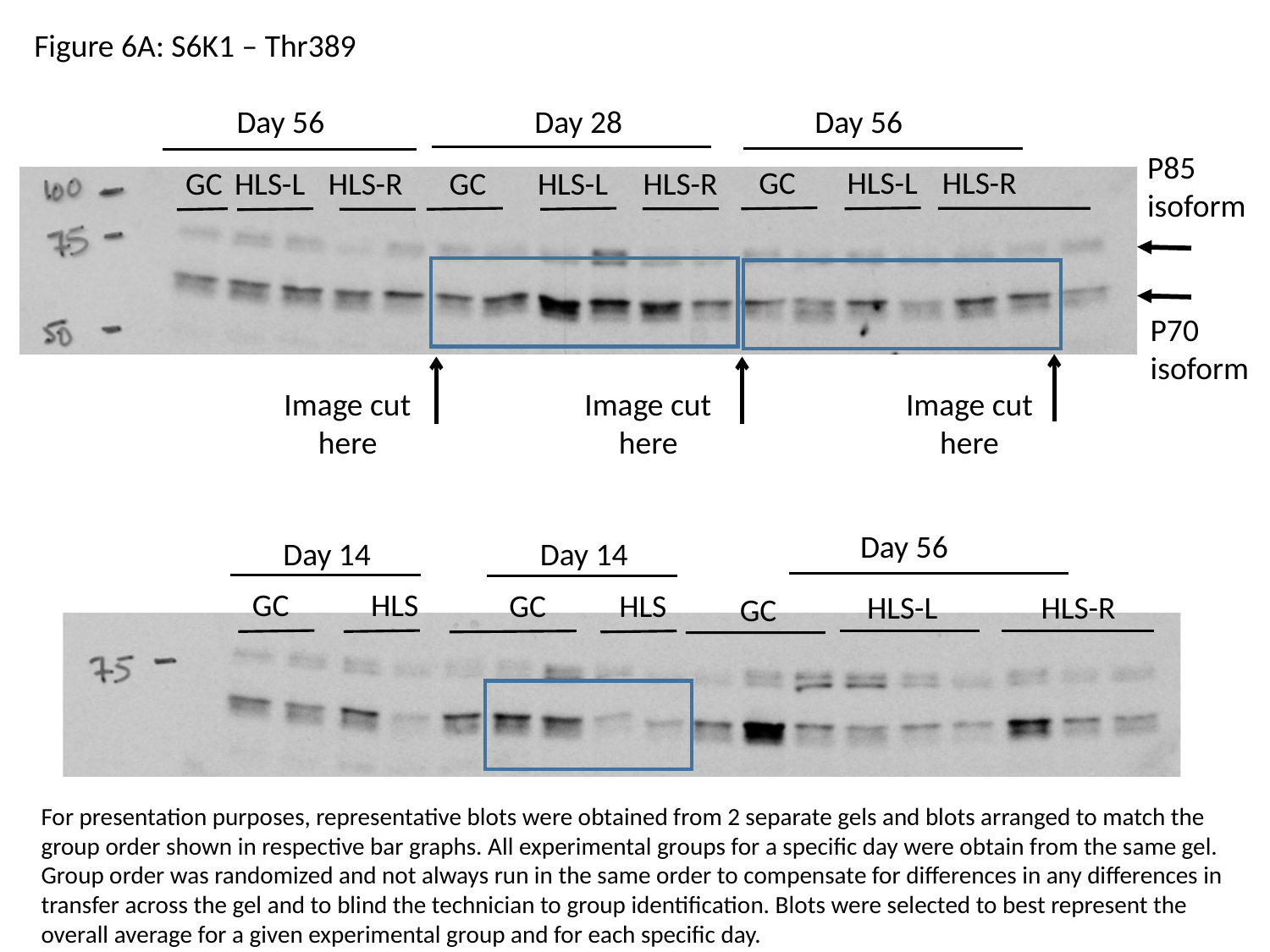

Figure 6A: S6K1 – Thr389
Day 56
Day 28
Day 56
P85
isoform
GC
HLS-L
HLS-R
GC
HLS-L
HLS-R
GC
HLS-L
HLS-R
P70
isoform
Image cut
here
Image cut
here
Image cut
here
Day 56
Day 14
Day 14
GC
HLS
GC
HLS
HLS-L
HLS-R
GC
For presentation purposes, representative blots were obtained from 2 separate gels and blots arranged to match the group order shown in respective bar graphs. All experimental groups for a specific day were obtain from the same gel. Group order was randomized and not always run in the same order to compensate for differences in any differences in transfer across the gel and to blind the technician to group identification. Blots were selected to best represent the overall average for a given experimental group and for each specific day.
